# Connecting the stimuli-responsive rheology of biopolymer hydrogels to underlying hydrogen-bonding interactions

**DOI:** 10.1101/2020.07.27.222802

**Authors:** Giulia Giubertoni, Federica Burla, Huib J. Bakker, Gijsje H. Koenderink

## Abstract

Many biopolymer hydrogels are environmentally responsive because they are held together by physical associations that depend on pH and temperature. Here we investigate how the pH and temperature response of the rheology of hyaluronan hydrogels is connected to the underlying molecular interactions. Hyaluronan is an essential structural biopolymer in the human body with many applications in biomedicine. Using two-dimensional infrared (2DIR) spectroscopy, we show that hyaluronan chains become connected by hydrogen bonds when the pH is changed from 7.0 to 2.5, and that the bond density at pH 2.5 is independent of temperature. Temperature-dependent rheology measurements show that due to this hydrogen bonding the stress relaxation at pH 2.5 is strongly slowed down in comparison to pH 7.0, consistent with the sticky reptation model of associative polymers. From the flow activation energy we conclude that each polymer is crosslinked by multiple (5-15) hydrogen bonds to others, causing slow macroscopic stress relaxation, despite the short time scale of breaking and reformation of each individual hydrogen bond. Our findings can aid the design of stimuli-responsive hydrogels with tailored viscoelastic properties for biomedical applications.

## INTRODUCTION

Hydrogels are an appealing class of polymer materials because of their responsiveness to environmental stimuli such as temperature, pH and ionic strength.^1^ Hydrogels derive this responsiveness from the presence of reversible associations between the constituent polymer chains, such as electrostatic, hydrogen-bonding, or hydrophobic associations or metal complexation^2^. Stimuli-responsive hydrogels have a wide range of applications, such as adhesives^3^, controlled drug delivery^4^, surrogate extracellular materials for tissue regeneration ^5^, and bio-inks for printing of soft tissues^6^.

For biomedical applications it is advantageous to create hydrogels from biopolymers that are naturally present in the human body, as these can capture both the mechanical and biological properties of the extracellular environment.^5^ A particularly interesting biopolymer in this context is hyaluronan, because of its ubiquity in human tissues and extracellular fluids such as the synovial fluid in joints and the vitreous humor of the eye.^7, 8^ Hyaluronan is a linear polyelectrolyte composed of repeating units of D-glucuronic acid and N-acetyl-D-glucosamine joined by glycosidic bonds.^9, 10^ In healthy tissues, hyaluronan has a very high molecular weight of up to 10 MDa, corresponding to chain lengths of tens of micrometers.^11^ Its length combined with its anionic nature enable hyaluronan to form soft hydrated viscoelastic fluids.^12–14^ These properties are at the basis of hyaluronan’s roles in the lubrication of joints^15^ and in tuning the transport properties and mechanical response of tissues such as skin, cartilage and brain^16, 17^. Hyaluronan’s mechanical properties together with its biochemical interactions regulate cell-matrix and cell-cell interactions, thus influencing cell-cell communication^18^, wound healing,^19^ and angiogenesis and development^20^. Accordingly, abnormalities in the expression level or molecular weight of hyaluronan brought about by aging or genetic disorders contribute to various diseases, such as inflammation^21^, multiple sclerosis^22^, cancer^23, 24^ and cartilage degeneration^25^.

Because of its central importance to human health and disease, there have been many studies of the material properties of isolated hyaluronan (reviewed in Ref.^14^). Like many other biological polyelectrolytes, hyaluronan is responsive to external stimuli, such as temperature, solution pH, salt concentration and multivalent counterions.^9, 26, 27^ Of these triggers, the solution pH has the most marked effect: in a narrow pH range centred around pH 2.5, hyaluronan solutions turn into soft solids that are stretchable.^28–31^ This special gel state is sometimes referred to as the *putty* state.^32^ We recently showed with two-dimensional infrared (2DIR) spectroscopy that this gelation process is caused by the pH-dependent formation of specific inter-chain hydrogen-bonds between the carboxyl and amide groups on adjacent chains (see Figure 1a).^33^

**Figure 1:**
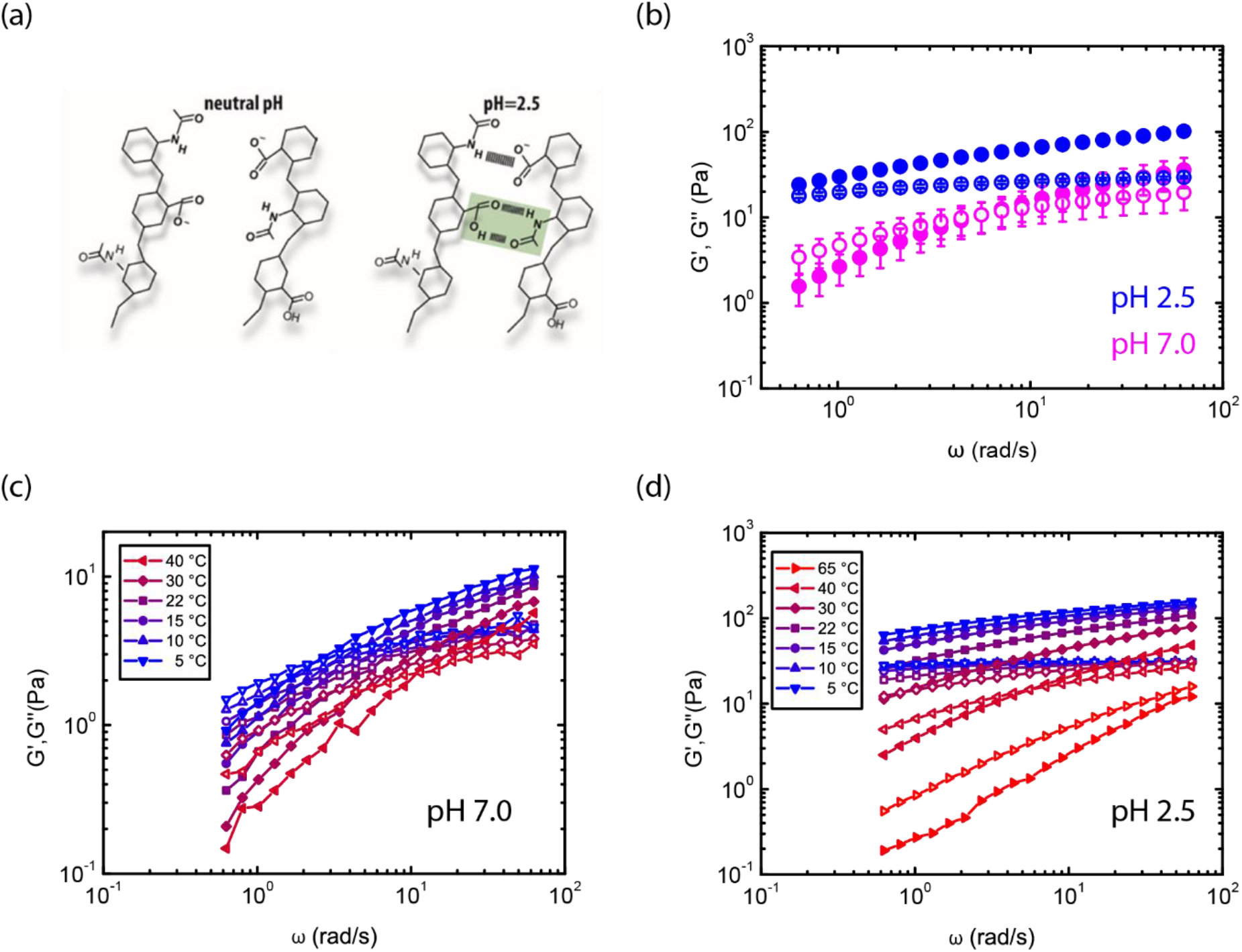
(a) Schematic showing the linear structure of hyaluronan, consisting of repeating disaccharides composed of N-acetyl-glucosamine and glucuronic-acid. At neutral pH (pH 7), the deprotonated carboxyl groups confer a net negative charge. At pH 2.5, the chains associate by the formation of hydrogen bonds between the carboxylic acid, the deprotonated carboxylate anion, and the amide groups of the hyaluronan polymers^39^. (b) Frequency spectra of the linear viscoelastic shear moduli of hyaluronan hydrogels (c = 10 mg/mL) at room temperature (T=22°C) at pH 7.0 (pink) and pH 2.5 (blue). Symbols denote the elastic (solid circles) and viscous (open circles) shear moduli. (c) Temperature dependence of the frequency spectra of the shear moduli of hyaluronan solutions at pH 7.0. (d) Corresponding frequency spectra at pH 2.5.

2DIR spectroscopy is uniquely capable of directly probing molecular interactions by measuring vibrational interactions using an intense femtosecond pump laser to excite the vibrations and a weaker probe beam to measure the induced vibrational responses. We found that at neutral pH all the carboxyl groups are deprotonated, and thus associative interactions between different polymer chains are prohibited by strong electrostatic repulsion. Hence, at neutral pH, the mechanical properties of hyaluronan solutions are dictated by chain entanglements. By lowering the pH to 2.5, close to the isoelectric point of hyaluronan^33^ of ~2.9, around 80% of the carboxyl groups are protonated. Protonation weakens the repulsive electrostatic forces between the chains and at the same time provides protonated carboxyl groups that participate in the formation of double hydrogen-bonds, together with amide groups and carboxylate anion groups (Figure 1a).

The material properties of responsive hydrogels can be tuned via the lifetime of the reversible crosslinks and their number density. There are several theoretical models for associative polymers that predict how the viscoelastic behavior depends on the crosslinker lifetime and density^34, 35^, but it remains challenging to understand the connection between macroscopic and molecular time scales. Hydrogen-bonds tend to be short-lived, forming and breaking on a timescale of a few to tens of picoseconds.^36^ Macroscopic stress relaxation in hyaluronan gels at pH 2.5 occurs on much longer time scales, ranging up to minutes. Here, we study the impact of temperature and pH on the rheology of hyaluronan hydrogels and on the underlying hydrogen-bond-mediated crosslinking by combining rheology and 2DIR spectroscopy. We compare hyaluronan at neutral pH, where the chains interact by excluded volume interactions (i.e. topological entanglements) only^26, 37, 38^, and at pH 2.5, where the chains additionally interact by transient hydrogen-bond associations^39^. We use time-temperature superposition rheology to measure the time scale and activation energy for network flow, and interpret these findings in the context of the sticky reptation model^34, 35^ previously applied to entangled synthetic polymers interacting by thermoreversible cross-linking^40, 41^. We use 2DIR spectroscopy to directly measure the density of interchain hydrogen-bonds as a function of temperature.

## EXPERIMENTAL SECTION

### Materials and sample preparation

Hyaluronan sodium salt in powder form obtained by fermentation of *Streptococcus equii* with a nominal molecular weight of 1.5-1.8 MDa was obtained from Sigma Aldrich. This molecular weight corresponds to a chain length of 3-4 μm and about 3750 disaccharides (considering that the disaccharide units have a molecular weight of 400 daltons and a length of ~1 nm [ref. ^42^]). For rheological measurements, solutions of HMW hyaluronan with concentrations between 5-20 mg/mL were prepared by dissolving accurately weighed amounts of hyaluronic acid in an aqueous solution containing 0.15 M NaCl (Sigma Aldrich) and HCl (Sigma Aldrich) ranging from 0 to 40 mM. The solution pH was measured with a pH meter equipped with a microelectrode (Hanna Instruments, Germany). The samples were left to equilibrate at room temperature for at least 24 hours before measuring. Samples were stored for at most 1 week at a temperature of 4°C. For the spectroscopic experiments, solutions of HMW hyaluronan in the putty state were prepared by dissolving 20 mg/ml hyaluronan sodium salt in heavy water at a DCl concentration of 40 mM, which was previously determined to be the correct concentration to obtain the pH=2.5 putty state conditions.^39^ Note that the pD value of a D_2_O solution can be converted to the pH value of an H_2_O solution of similar acidity using pH = pD*0.929.^43^

### Time-temperature superposition rheology experiments

We measured frequency spectra of the linear viscoelastic shear moduli as a function of temperature with a stress-controlled MCR 501 rheometer (Anton Paar, Austria), using a plate-plate measuring geometry with a 40 mm diameter and 100 μm gap. The plate temperature was controlled by a Peltier plate connected to the lower plate and a hood controlling the temperature of the upper plate. Samples were loaded at room temperature (22°C) and equilibrated for 10 minutes before starting the experiments. Small amplitude (0.5%) oscillatory shear tests were performed at 20 frequencies logarithmically spaced between 0.1 and 10 Hz, at temperatures of 5°C, 10°C, 15°C, 22°C, 30°C and 40°C. We started at 22°C and first decreased the temperature to 15°C, 10°C, and 5°C, and then increased the temperature to 30°C and 40°C. The temperature was adjusted between each frequency sweep at the fastest rate allowed by the rheometer (around 2°C/s), and after reaching the desired temperature, we waited for equilibration by waiting for the elastic and loss moduli to saturate, which always occurred within two minutes. We verified that there was no hysteresis in the viscoelastic moduli upon cooling back to 22°C until we raised the temperature to 60°C (Fig. S1), consistent with prior work,^44^ that showed that hyaluronan is stable up to 50°C and is hydrolysed above 90°C on a timescale of ~1 hour. Time-temperature superposition analysis of the frequency (ω) spectra was performed by determining the shift factor *a*_T_(*T*) required to overlap both the elastic modulus *G*’(*ω*) and the viscous shear modulus *G*”(*ω*) with their respective reference curves measured at the reference temperature *T*_0_ = 295 K (22°C). The temperature dependence of the shift factors *a*_T_(*T*) was fitted to an Arrhenius equation to obtain the activation enthalpy *E*_a_ for stress relaxation:

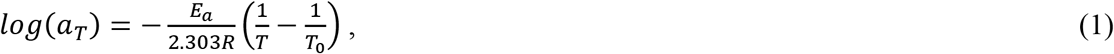

where *R* is the universal gas constant. The activation energy in units of *k*_B_*T* is obtained by dividing by Avogadro’s number, where k_B_ is Boltzmann’s constant and *T* is the absolute temperature. At each temperature we performed experiments on three independently prepared samples.

### Linear infrared spectroscopy (FTIR)

We measured Fourier Transform Infrared (FTIR) spectra of the sampled with a Bruker Vertex 80v FTIR spectrometer equipped with a liquid-nitrogen-cooled-mercury-cadmium-telluride (MCT) detector. The spectra were recorded under nitrogen atmosphere at a wavenumber resolution of 3 cm^−1^. We averaged 100 scans for every spectrum. In all measurements we used a standard sample cell with a path length of 100 μm. The temperature-dependent FTIR spectra were measured with the aid of a Peltier-cooled temperature cell (Mid-IR Falcon, Pike technologies). The temperature was ramped from 20°C to 40°C at a rate of 1°C/min. The spectra were corrected for the absorption of the D_2_O solvent background at the same temperature. These background spectra were measured using the same ramping parameters. To determine if the ratio between the two protonation states of the carboxyl group depends on temperature, we fitted the linear infrared spectra measured at different temperatures using a global fitting procedure. This fit is based on the minimization of the square error:

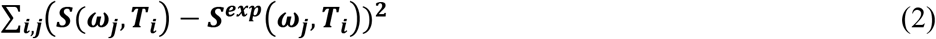

where *S* is the fitted spectrum, *S*^exp^ the experimental spectrum, *ω*_j_ the frequency, and *T*_*i*_ the temperature. Assuming that the vibrational bands are Gaussian shaped, we expect that

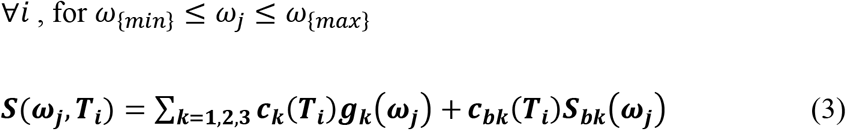

where the *g*_k_ represent the three Gaussians that describe the absorption bands of the anti-symmetric stretching mode of the carboxylate anion (COO^−^_ant_), the amide I mode (AM.I) and the carbonyl stretching mode of the protonated carboxyl group (C=O), respectively, and *c*_k_ are the Gaussians amplitudes. *S*_*bk*_(*ω*_*j*_) represents the spectrum used for background correction. Based on previous work^39^, the Gaussians were centered at 1609 cm^−1^, 1634 cm^−1^ and 1725 cm^−1^ for the absorption bands of the COO^−^_ant_, AM.I and C=O vibrations, respectively. To account for the fact that increasing the temperature leads to a slight shift of the absorption bands^45^, the center frequencies were allowed to shift by a maximum of 1 cm^−1^ when fitting the linear spectrum at 40°C. The widths of the three bands are global parameters in the fit, meaning that they are fixed at all studied temperatures, and only the amplitudes of the three bands are allowed to be different at each temperature. We can then find the areas of the COO^−^_ant_, AM.I and C=O absorption bands at each temperature, and compare their ratio at different temperatures.

### Two-dimensional infrared spectroscopy (2DIR)

2DIR measurements were performed using a home-built setup described elsewhere.^46^ We excite the amide and carboxyl vibrations with a pair of strong femtosecond mid-infrared pulses centered at 1670 cm^−1^ with a bandwidth of 200 cm^−1^ (~100 fs, 4 μJ per pulse). This excitation induces transient absorption changes that we probe with a weaker (0.35 μJ) single femtosecond probing pulse that is delayed by a waiting time *T*_w_. After propagation through the sample, the probe pulse was sent into an infrared spectrograph and detected with an infrared mercury-cadmium-telluride (MCT) detector array, thus yielding the transient absorption spectrum as a function of the probe frequency. The dependence of the transient absorption spectrum on the excitation frequency was determined by measuring transient spectra for many different delay times *τ* between the two excitation pulses. By Fourier transformation of these spectra, we obtain the dependence of the transient absorption spectrum on the excitation frequency. By plotting the transient absorption spectrum as a function of the excitation and the probing frequency, we obtain 2D-IR transient absorption spectrum for each delay time *T*_*w*_ between the excitation and probing pulses. The temperature of the sample was kept constant during the experiment with a Peltier element with an active feedback loop.

## RESULTS AND DISCUSSION

### pH dependence of hyaluronan hydrogel rheology

To study the impact of pH-dependent crosslinking on hyaluronan hydrogel rheology, we compare solutions of hyaluronan polymers at pH 2.5 and pH 7.0 (see Figure 1a). These pH values were chosen because they represent two distinct regimes of hyaluronan chain interactions. At pH 7.0, previous rheology^37^ and self-diffusion^26^ measurements showed that hyaluronan chains interact by excluded volume interactions only, provided that the ionic strength of the solution is sufficiently high to screen the electrostatic repulsions of the chains. At pH 2.5, high molecular weight hyaluronan forms soft and stretchable gels that are referred to as *putty*.^32, 47^ We recently showed with 2D-IR spectroscopy measurements that this gel formation is driven by the pH-dependent formation of hydrogen bonds involving carboxylic acid, the deprotonated carboxylate anion, and the amide groups of adjacent hyaluronan chains (Fig. 1a).^39^ We performed the rheology experiments on samples with a polymer concentration of 10 mg/ml, in the *semidilute entangled* concentration regime. In this regime, the polymers experience transient topological constraints as a result of their high density.^14^ The entangled regime sets in at the entanglement concentration ce, which is generally a factor 2-5 higher than the overlap concentration c^*^ where the polymer coils first touch.^48^ For hyaluronan with a molecular weight of 1.5-1.8 MDa, *c*\* ≈ 2 mg/ml.^49, 50^ We characterized the time-dependent rheology of the samples by performing frequency sweeps using strain oscillations with a small (0.5%) amplitude.

As shown in Fig. 1b, the frequency spectra show a transition from a solid response (storage shear modulus *G*’ larger than the loss modulus *G*”) at high frequencies to a fluid response (*G*’ <*G*”) at low frequencies. This behaviour is characteristic for entangled polymer solutions where chains diffuse by reptation, i.e. highly restricted diffusion due to entanglements with surrounding chains^48^. Entangled polymer solutions generally display a Maxwell-type behaviour with a rubber-like plateau that persists down to a frequency *ω*≈1/*τ*, and a terminal (*ω*<1/*τ*) regime where the moduli tend towards *G*”∝*ω*^2^ and *G*’ ∝*ω*^1^ scaling^48^. The main effects of lowering the pH are first a down-shift of the frequency where *G*’ and *G*’’ cross-over, which marks the transition from the rubber plateau to the terminal regime (from 5 rad/s at pH 7.0 to ~0.2 rad/s at pH 2.5), and second an approximately 10-fold increase of the plateau modulus (from ~20 Pa at pH 7.0 to ~200 Pa at pH 2.5). These changes are as expected for entangled polymers that experience transient crosslinking interactions, as formulated by the *sticky reptation* model.^34, 35^ This model predicts that sticky interactions slow down reptation by adding an additional friction on the chains that increases linearly with the (effective) bond lifetime and quadratically with the number of stickers per chain. The model further predicts that transient crosslinks enhance the shear modulus *G*_0_ in the rubbery plateau. *G*_0_ is the sum of the free energy of the strands in the material, *G*_0_=*v*_x_k_B_T, where *v*_x_ =*c*N_A_/*M* is the number density of elastically active network strands, *N*_A_ is Avogadro’s number, and *M* is the chain molecular weight. Assuming that the number densities of constraints provided by entanglements and crosslinks add up, the data indicate that reducing the pH from 7.0 to 2.5 creates a 10-fold increase in the density of elastic constraints.

### Temperature dependence of hyaluronan hydrogel rheology

Our findings suggest that the rheology of hyaluronan hydrogels is controlled by a single time scale, which is modulated by pH-dependent chain interactions. To test this hypothesis, we performed measurements at different temperatures, since temperature is known to change the time scale of hydrogen-bond breaking and reformation.^51^ As shown in Figure 1c, the shear moduli both at neutral (pH 7.0) and at acidic (pH 2.5) hyaluronan solutions strongly depend on temperature. Increasing the temperature lowers the magnitude of the shear moduli and increases the characteristic frequency at which the rubber plateau crosses over to the terminal regime, consistent with faster stress relaxation. We note that the hyaluronan hydrogels were thermoreversible: there was no hysteresis in the rheology when the temperature was shifted back up or back down (Fig. S1 in the Supplementary Information).

To test whether a single interaction mechanism can indeed account for the observed temperature dependence, we performed a time-temperature superposition analysis using 22^°^C as the reference temperature. The idea behind this analysis is that the stress relaxation time according to the sticky reptation model is proportional to the (effective) bond life time and should therefore be exponentially dependent on temperature.^52^ As shown in Figure 2, we can indeed construct master curves for both the pH 7.0 data set (Fig. 2a) and the pH 2.5 data set (Fig. 2b) by in each case shifting the data along the frequency axis by a temperature-dependent shift factor aT. The fact that the same horizontal shift factors apply for *G*’ and *G*” indicates that hyaluronan gels are thermorheologically simple materials^47^ for which a single temperature-dependent interaction scale determines the rheology. The fact that we only need to apply a horizontal (time) shift factor to superpose the frequency-dependent *G*’/*G*” curves shows that only the time scale, not the equilibrium density, of crosslinks changes with temperature. The master curves clearly show Maxwell-type behavior with a rubbery plateau at the higher frequencies and a crossover to a terminal relaxation regime at lower frequencies. At pH 7.0, we do not reach sufficiently low frequencies to confirm whether the moduli at low frequencies tend towards the *G*’’ ~ ω^1^ and *G*’ ~ ω^2^ dependencies expected for a Maxwell fluid with a single relaxation time. At pH 2.5, we do reach lower normalized frequencies, but the low-frequency scaling of *G*’ seems somewhat shallower than the *G*’ ~ ω^2^ scaling of the Maxwell model. A likely explanation is the size polydispersity of the hyaluronan, which is known to cause broadening of the frequency spectrum of entangled polymers^48^, especially in the case of sticky polymers^41^.

**Figure 2.**
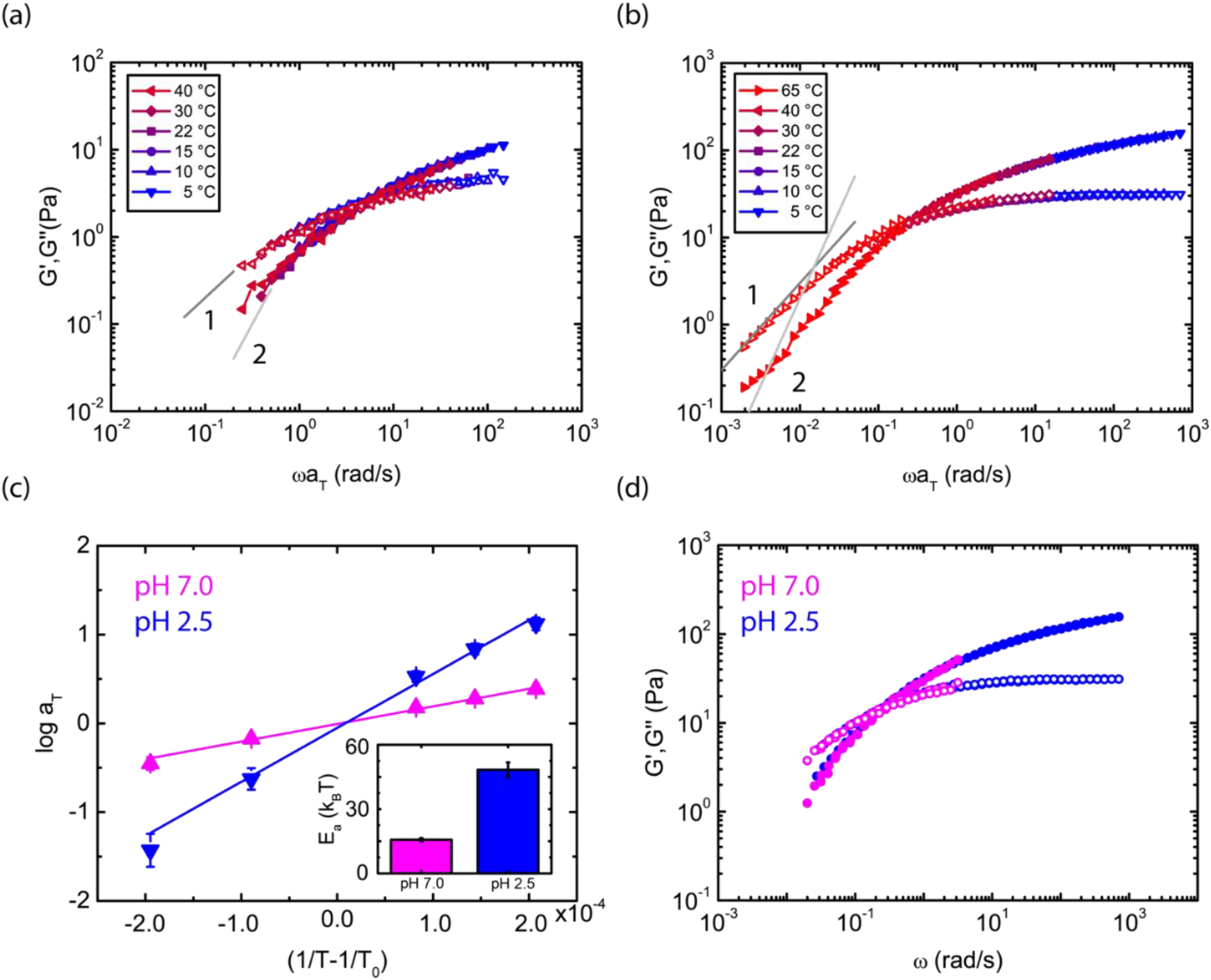
Time-temperature superposition (TTS) rheology of hyaluronan solution (10 mg/ml). (a) Frequency sweeps at pH 7.0 obtained at different temperatures are shifted along the frequency axis onto the measurement at the reference temperature of 22°C by multiplication with a shift factor a_T_. (b) Corresponding rescaled frequency sweeps at pH 2.5. The grey lines in (a,b) indicate the expected terminal relaxation *G*”∝*ω*^1^ and *G*’∝*ω*^2^ for a Maxwell fluid. (c) Arrhenius plot showing the shift factors used for constructing the time-temperature superposition data as a function of the inverse of the temperature relative to the reference temperature. Lines show linear fits whose slopes represent the flow activation energy divided by 2.3R. Inset: Activation energies retrieved from the Arrhenius fits, expressed in units of thermal energy k_B_T. (d) Universal master curve for hyaluronan hydrogel rheology obtained by rescaling all frequency sweeps gathered at different temperatures (5 and 40°C, not color-coded) and pH values (2.5 and 7.0, see legend) onto a reference set measured at pH 2.5 and at 22°C (blue triangles). We first shifted the horizontal axis, then the vertical axis to obtain overlap.

To retrieve the pH-dependence of the flow activation energy, we constructed Arrhenius plots of the shift factors *a*_T_ as a function of the inverse temperature relative to the reference temperature *T*_0_=22°C, both for pH 7.0 and pH 2.5.^53^ As shown in Figure 2b, we obtained straight lines at both pH values, indicating that the relaxation time is controlled by the thermally activated dissociation of bonds between chains. As shown in Fig. 2c, the activation energy at pH 2.5 (taken from the slope of the Arrhenius plots) is much higher than at pH 7.0, consistent with the expectation from the sticky reptation model that the formation of hydrogen bonds between the chains enhances the friction on the chains. In the entangled state at pH 7.0, the activation energy is around 12 k_B_T (or 30 kJ/mol), consistent with prior findings.^50^ We note that the earlier work was carried out at 50 mg/mL, a much larger concentration than studied here (10 mg/ml), suggesting little effect of concentration on the activation energy. We also find little dependence of the activation energy on hyaluronan concentration at pH 7.0 when we extend the concentration range down to 5 mg/ml (see Supplementary Figure S2). For the putty state at pH 2.5, the activation energy is much higher and amounts to ~50 k_B_T (or 115 kJ/mol). We measured similar activation energies at other hyaluronan concentrations (5-20 mg/ml) (see Supplementary Figure S2). The much higher flow activation energy at pH 2.5 compared to pH 7.0 is consistent with the additional presence of transient associations between the chains.

Until now, we rescaled the data sets for different pH values onto separate master curves. To test if a single constitutive law describes all data, independent of pH and temperature, we finally rescaled all curves onto a single master curve. This idea was inspired by recent studies showing that time-temperature superposition can be extended to other environmental parameters that control the relaxation times of noncovalent gels such as ionic strength.^54, 55^ We can indeed rescale the *G*’ and *G*’’ curves measured at pH 7.0 and pH 2.5 onto a single universal set of master curves, using in addition to the horizontal shift factor *a*_T_ for the frequency axis also a vertical shift factor *b*_T_. The shift factor *b*_T_ is around 7, indicating a 7-fold higher density of elastic constraints at pH 2.5 compared to pH 7.0. The collapse onto a single set of master curves strongly suggests that the physical mechanism that controls the rheology is the same, regardless of temperature and pH.

### Spectroscopic measurement of the density of intermolecular interactions

Prior 2DIR spectroscopy measurements from our lab^39^ showed that the crosslinks between hyaluronan chains are formed by hydrogen bonds between the carboxylic acid, deprotonated carboxylate anion, and amide groups of adjacent hyaluronan polymers (Figure 1a). The crosslink density thus likely depends on the degree of protonation of the carboxyl groups, which could in principle change with temperature. To test whether the protonation state of hyaluronan changes with temperature, we measured linear infrared absorption spectra of a solution of hyaluronan dissolved in D_2_O at 20°C and 40°C (Fig. 3). The solution was prepared at a concentration of 40 mM of DCl in order to obtain a pH of 2.5 where hyaluronan is in the putty state, following the same protocol as previously described^39^. We observe three absorption bands, at 1633 cm^−1^, 1725 cm^−1^, and 1609 cm^−1^, which based on prior work^56^ we assign to, respectively, the amide I vibration (AM.I), the carbonyl stretching mode of the protonated carboxylic acid group (C=O), and the antisymmetric stretch vibration of the carboxylate group anion group (COO^−^_ant-_). The relative intensities of the C=O and COO^−^_ant_ bands directly reflect the protonation state of hyaluronan.^39^ We fit these bands following the procedure described in the Methods section and we determine the absorption band areas. We find that the fraction of deprotonated carboxyl groups (− COO^−^) is the same at 20°C and at 40°C. From the acid-base equilibrium of hyaluronan, we calculate that ~20/30% of the −COO- is deprotonated at pH 2.5.

**Figure 3.**
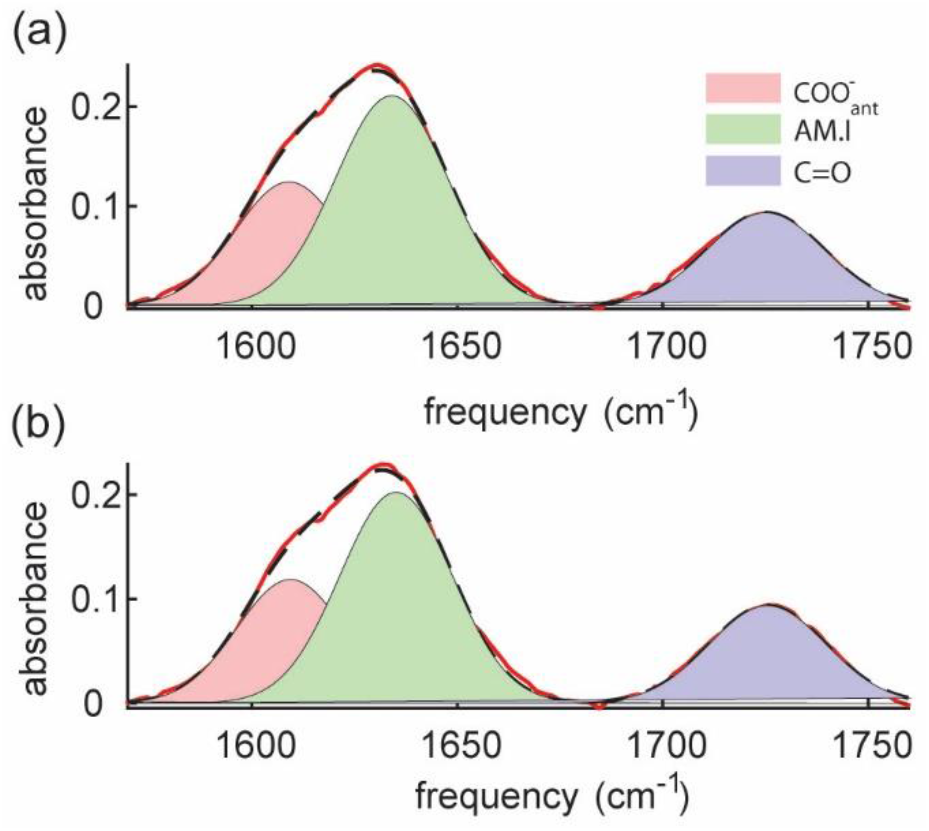
(a) and (b) linear infrared spectra for a solution of hyaluronan of 20 mg/ml at pH=2.5 at two temperatures of 20°C and 40°C, respectively. Red solid lines represent the experimental data, while dashed black lines show the fits. By calculating the areas of the absorption bands, we find that the ratio between −COO- and −COOH is constant (~1.1).

To measure the density of hydrogen-bonds as a function of temperature, we performed 2D-IR spectroscopy measurements on hyaluronan solutions in D_2_O at pH=2.5 at several temperatures between 5°C and 40°C (Fig. 4). We excited the amide and carboxyl vibrations with a strong ~100 femtosecond infrared pulse pair centered at 1670 cm^−1^. We probed the resulting transient absorption changes with a weaker (0.35 μJ) single femtosecond probing pulse that was delayed by a waiting time T_w_ (Figure 4a). Figure 4b and c show the isotropic 2DIR spectra at the two temperature extremes of 5°C and 40°C. In both spectra, we observe on the diagonal four distinct spectral signatures. The blue-coloured signatures represent negative absorption changes (which we refer to as *bleach*) and originate from a reduction of the absorption at the amide I and carboxyl carbonyl frequencies. This reduced absorption is due to the depletion of the fundamental transition (*v*=0 to *v* =1) and stimulated emission (*v* =1 to *v* =0) of the excited amide I and carbonyl vibrations. At somewhat lower probing frequencies, we observe red-colored spectral signatures, which represent positive absorption changes. These positive absorption changes can be assigned to excited state absorption or *esa* and arise because of the vibrational transition from *v* =1 to *v* = 2.^57^ The two pairs of diagonal peaks are obtained by exciting at 1633 and 1725 cm^−1^, corresponding to the vibrational frequencies of the amide I and C=O absorption bands, respectively. The lower-frequency diagonal peak shows a shoulder at 1610 cm^−1^, which we assign to the anti-symmetric stretch vibration of the carboxylate anion.

**Figure 4.**
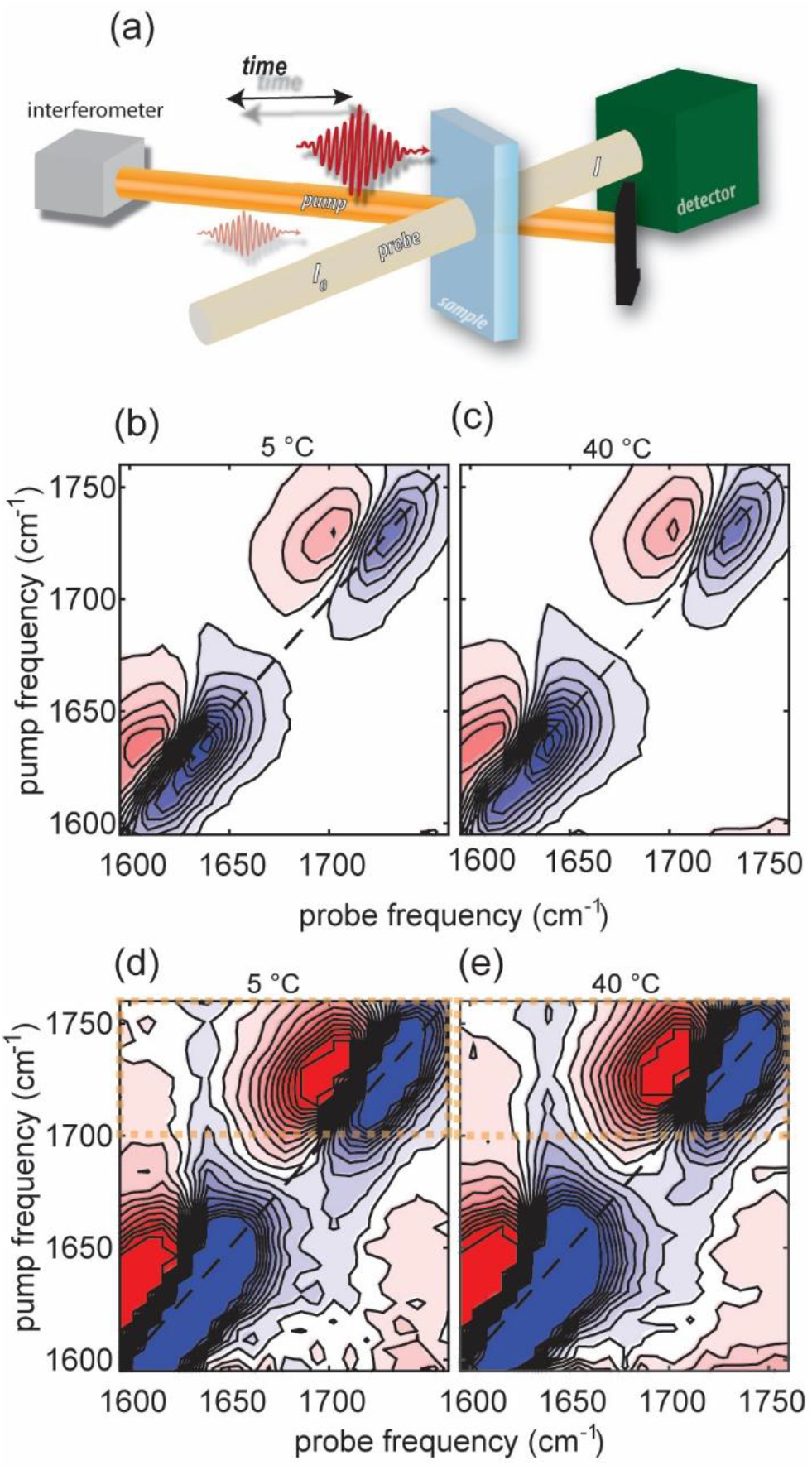
(a) Schematic of the 2DIR experiment (see main text for details). (b-c) 2DIR spectra of hyaluronan (20 mg/ml) at pH=2.5 at a time delay T_w_=0.3 ps at two temperatures, namely 5°C (b) and 40°C (c). (d,e) Same spectra with color-bar fivefold magnified, to show the presence of off-diagonal peaks in the frequency regions indicated by the orange dashed rectangles.

When we multiply the 2DIR spectra by a factor of 5, we can also observe off-diagonal spectral signatures in the 2DIR spectrum at both temperatures (Fig. 4d,e). The off-diagonal peaks are ten-fold weaker than the diagonal signals. In the top-left off-diagonal region (exciting around 1725 cm^−1^ and probing around 1630 cm^−1^), which is indicated by the orange-colored rectangle, we observe a 2DIR absorption signal associated with the excitation of the carbonyl vibration and probing of the amide I absorption. This off-diagonal feature is commonly referred to as a downhill cross-peak. Fig. 5a reports the 2DIR signals obtained by averaging over the excitation frequencies between 1700 and 1760 cm^−1^ (orange rectangle in Fig. 4d-e). At each temperature, the 2DIR signal is normalized to the maximum intensity of the carbonyl bleach. In Fig. 5a we observe again the bleach and the esa of the carbonyl around 1700 cm^−1^, and a negative 2DIR signal near 1635 cm^−1^, which is the absorption frequency of the amide I vibration. The cross-peak signal between the carbonyl stretching and the amide I vibrations reveals the existence of molecular coupling between these groups at pH=2.5, and shows that a fraction of the carboxyl and the amide I groups are connected by a hydrogen bond.

**Figure 5.**
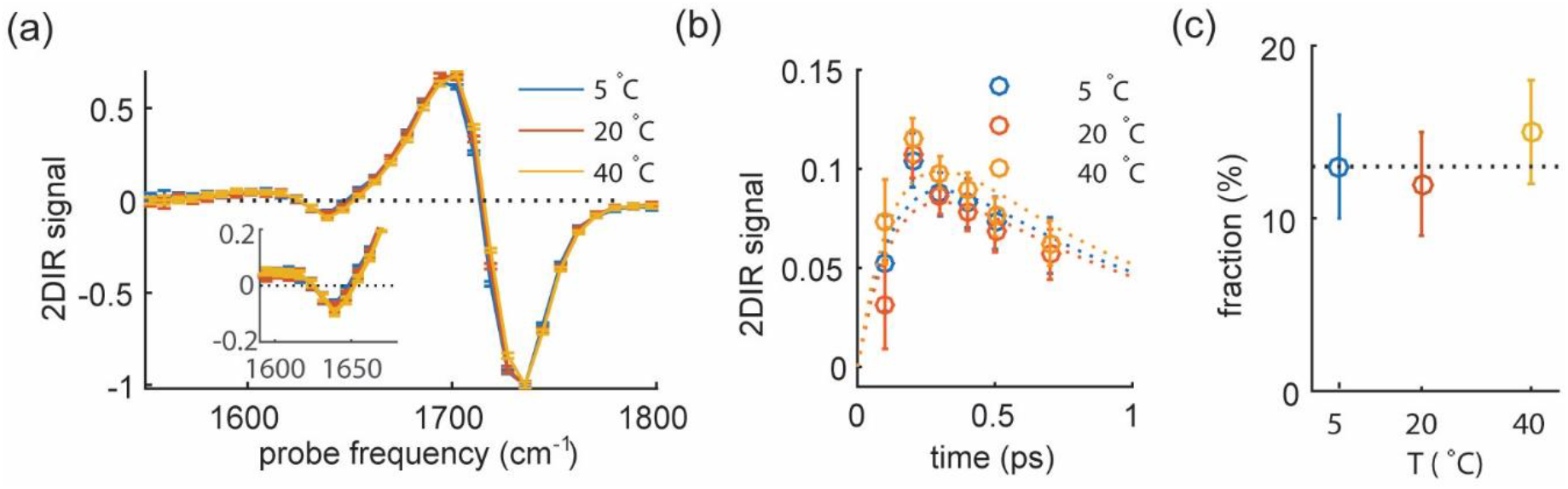
Temperature dependence of 2DIR spectral features for hyaluronan hydrogels at pH=2.5. (a) 2DIR signals at three different temperatures obtained by averaging over the pump region between 1700 and 1760 cm^−1^. Inset: detail of the 2DIR signal in the absorption frequency of the amide I vibration showing the cross-peak signal is independent of temperature. (b) Cross-peak decay traces at 5, 20 and 40°C. Dashed lines are fits to the relaxation model described in the main text. (c) The fraction of hydrogen-bonded carboxylic acid groups extracted from the 2DIR data is independent of temperature.

To extract the fraction of hydrogen-bonded carboxyl and amide I groups, we consider the delay-time traces of the cross-peak signals shown in Fig. 5b, obtained by fitting the 2DIR signals reported in Fig. 5a (see Supplementary Materials section in SI for details). The three delay-time traces are very similar, not showing a significant change upon changing the temperature. In order to extract the fraction of the hydrogen-bonded carboxyl and amide I groups at different temperatures, we analyzed the delay-time traces. In an earlier 2DIR study of hyaluronan^39^, we showed that upon excitation, the carbonyl stretching mode can relax following two different relaxation paths.

A fraction of the excited C=O vibrations relaxes by fast energy transfer to the amide I vibration with a time constant *T*_ent_ (~0.150 ps). This fast relaxation is enabled by the formation of the intermolecular hydrogen bonds. This energy transfer process leads to excitation of the amide I vibration and thus to a fast rise of the cross-peak signal. Subsequently, the amide I mode relaxes back to the ground state with a time constant *T*_1AM.I_ (~0.65 ps). Another fraction of the excited C=O groups is not hydrogen-bonded to an amide group and relaxes to low-frequency modes of the solvent or the hyaluronan polymer chain. This process is much slower and occurs with a time constant *T*_1COOD_ (~ 0.7 ps). Once populated, the low frequency modes affect the absorption spectrum of the amide I vibration and also generate a 2DIR cross-peak signal. This latter contribution to the signal decays with a time constant *T*_1LFM_, which is the relaxation time of the low frequency modes. In our previous work^39^, we found that *T*_1LFM_ is ~3 ps, which is much slower than the decay of the amide I and C=O vibrations. Because the two relaxation mechanisms that determine the cross-peak dynamics occur on quite different time scales, they can be distinguished in a fit of the cross-peak delay-time traces with the following expression:

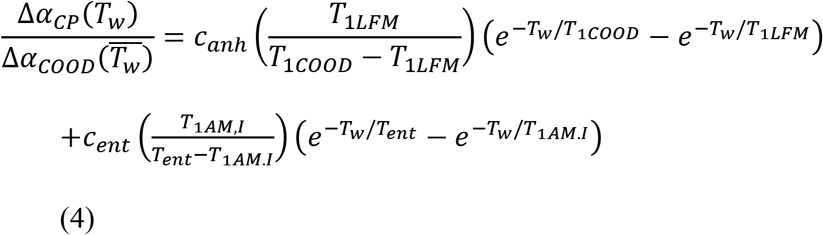

where 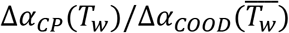 is the intensity of the cross-peak normalized to the intensity of the C=O vibration at early time delay and 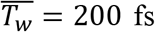. The parameters *c*_anh_ and *c*_ent_ represent the cross-peak amplitudes associated with the anharmonic coupling and direct energy transfer mechanisms, respectively, and are the only free parameters. Because of the limited amount of data points at long time delay, the value of *c*_anh_ is constrained to be a least 0.02, which is the minimum value expected for this parameter based on our previous work (*c*_anh_=0.025±0.005). Hence, we extract the amplitude of the fast relaxation component due to energy transfer, which can be directly linked to the fraction of protonated carboxylic acid bonded to the amide groups. We find that around 15% of the carbonyl groups are hydrogen-bonded to the amide groups at pH=2.5 (Fig. 5C). This fraction does not change in the temperature interval between 5 and 40°C.

### Connection between rheology and 2DIR

The 2DIR measurements show that the fraction of carboxyl groups that are engaged in a direct hydrogen-bond with an amide group does not change with temperature between 5 and 40°C. This finding is consistent with the time-temperature superposition analysis of the rheology data, which showed that the density of stickers that impede stress relaxation also does not change with temperature. These findings indicate that the breaking of an interchain hydrogen bond between a carboxyl group and an amide group results in a state that has about the same overall free energy as the intact hydrogen bond (see schematic in Figure 6). This other state is probably formed by hydrogen bonds between the carboxyl and the amide groups and solvent water molecules. As a result, a change in temperature does not affect the number of hydrogen-bonds that are formed between the polymer chains. However, the kinetics do change with temperatures, as the breaking and reformation involves an energy barrier *E*_A,hb_ that will be of the order of the binding energy of a single hydrogen bond, which is typically ~400-1400 cm^−1^ or 2-7 k_B_T (T=298 K).^58, 59^

**Figure 6.**
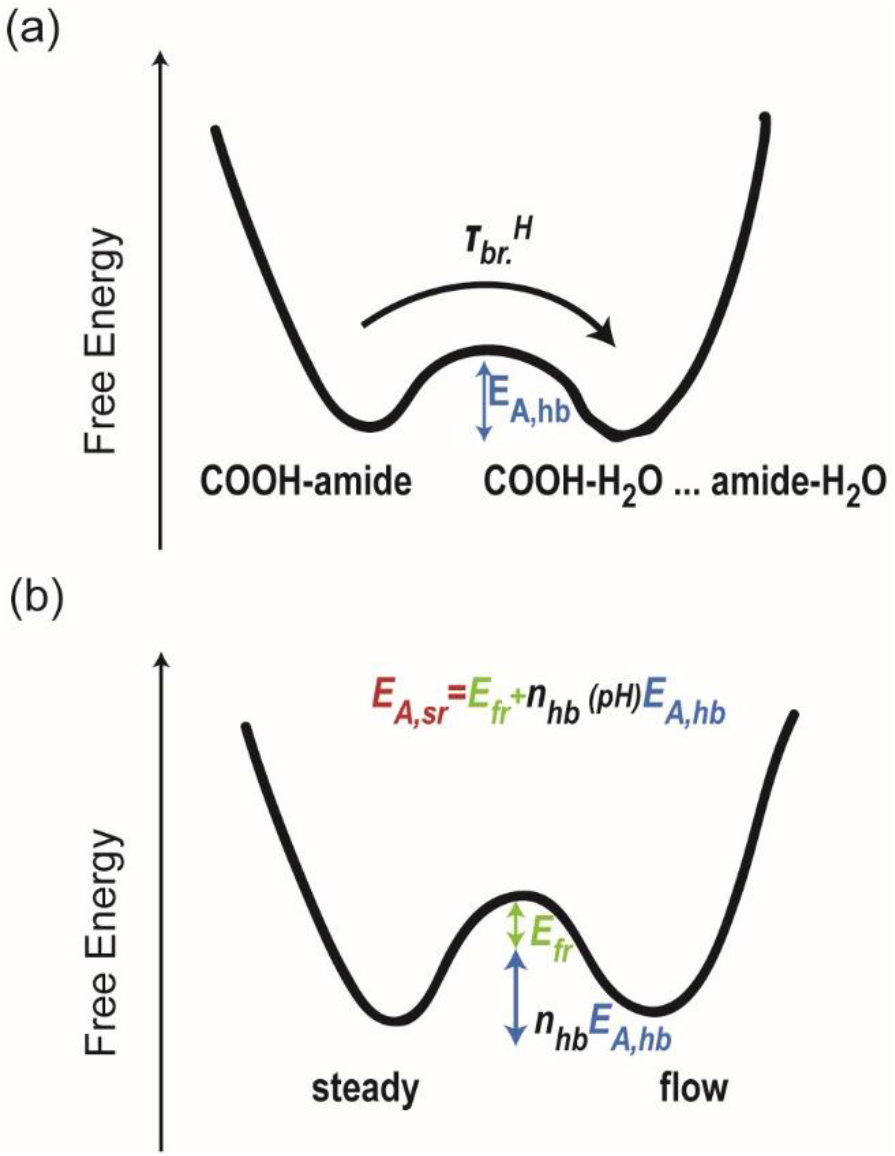
a) Schematic representation of the breaking of an hydrogen-bonded complex. We refer to the breakage time of the hydrogen-bonds as 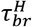. The activation energy necessary to break the hydrogen-bonded complex is defined as *E*_*A,hb*_. b) Schematic representation of the network dynamics. The flow activation energy *E*_*A,sr*_ corresponds to the breaking of *n*_hb_ hydrogen bonds (stickers).

To understand the acceleration of the network dynamics with increasing temperature, we need to find the relation between the characteristic lifetime of a single hydrogen bond *τ* and the characteristic time scale of the stress-relaxation of the hyaluronan solution, as expressed in the frequency dependence of the shear moduli. The typical lifetime 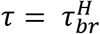 of a single hydrogen-bond is on the order of a few to tens of picoseconds.^36^ This large disparity in time scales between single hydrogen bond dynamics and sticker dynamics can be understood from the fact that multiple hydrogen bonds *n*_hb_ need to be broken at the same time to allow for chain displacement, and thus flow of the network. In addition, such a mesoscopic rearrangement involves a frictional barrier *E*_*fr*_ due to chain entanglements and viscous drag with the solvent. Hence, the activation energy necessary for the network to flow will be larger than the activation energy of breaking a single hydrogen bond. Assuming a similar pre-exponential (Arrhenius) prefactor for the breaking of hydrogen bonds and the friction, the total activation energy for stress relaxation can be written as:

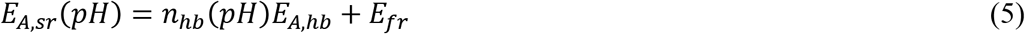

At pH=7 we can assume that there are no hydrogen bonds (n_hb_=0) formed between the hyaluronan polymer chains based on earlier work showing that excluded volume interactions alone are enough to explain the concentration-dependent rheology.^37^ Thus, the activation energy at pH 7 will be equal to the frictional barrier energy *E*_fr_. At pH=2.5, hydrogen bonds are formed between the hyaluronan polymer chains and their density is large enough to cause the formation of an elastic hydrogel. By subtracting the frictional energy *E*_fr_ from the activation energy measured at pH=2.5, *E*_A,sr_, we find that the energy barrier due to hydrogen-bonds at pH 2.5 is around 85 kJ/mol (~ 38 k_B_T). This residual activation energy corresponds to *n*_hb_=5-15 hydrogen bonds, which thus need to be broken in order for the hyaluronan chain to diffuse at pH=2.5.

Finally, it should be noted that we have not considered the possible effect of temperature on the frictional energy barrier *E*_fr_. This energy barrier will depend on several parameters, such as the intramolecular hydrogen-bonds that stiffen hyaluronan and the mobility of the solvent molecules. Increased chain flexibility and water mobility could potentially lower *E*_fr_ as the temperature increases^60, 61^. However, the fact that the temperature dependence of the shift factor aT can be well described with an Arrhenius expression suggests that the value of *E*_fr_ is quite insensitive to temperature changes within the range (5-40°C) we studied.

## CONCLUSIONS

We used rheological and two-dimensional IR spectroscopy measurements to measure the effect of temperature and pH on the mechanical properties and underlying molecular interactions of entangled hyaluronan polymers. The rheology results showed that changing the pH from 7.0 to 2.5 introduces associative interactions that strongly slow down stress relaxation. The 2DIR results showed that these associative interactions are mediated by hydrogen bonds whose equilibrium density does not depend on the temperature (from 5° C to 40°C) but only on pH. Hence, the equilibrium structure of the hyaluronan polymer network does not change with temperature. The kinetics of the polymer network does show a strong dependence on temperature. By performing measurements over an extended temperature range (5-40°C), we could show that the stress relaxation time displays an Arrhenius-type dependence on temperature with a flow activation energy of 12 k_B_T (⁓30 kJ/mol) at pH 7.0 and 50 k_B_T (⁓115 kJ/mol) at pH 2.5. The activation energy at pH 7.0 can be assigned to a frictional energy barrier associated with the motion of the polymer chains along each other and the displacement of water molecules. The much higher activation energy at pH=2.5 indicates that under these conditions, the simultaneous breaking of 5-15 hydrogen bonds is required for chains to move and, thus, for the network to flow. Our findings show that hyaluronan forms thermoreversible gels whose macroscopic properties can be tuned by pH and temperature.

In the context of human physiology, understanding how the mechanical properties of extracellular glycosaminoglycans such as hyaluronan are affected by pH is relevant since many diseases, like cancer and inflammation, are associated with acidosis, a decrease of the extracellular pH ^62–64^. Even though the average pH values in this situation are reported to be of order 5.8 [ref.^65^], potentially lower pH values might be reached locally and transiently, which may strongly affect the dynamics of the hyaluronan polymer network. Moreover, pH-responsive hydrogels may potentially serve for targeted release of anticancer drugs^66^. In the context of materials science, it has become clear in recent years that noncovalently bonded hydrogels have many desirable properties such as stimuli-responsiveness, toughness, and an inherent capacity for self-healing and recovery after mechanical damage. Such materials have possible applications for shape memory, self-healing and adhesive materials^1^. Hyaluronan is biocompatible and already used for a broad range of biomedical applications, from drug delivery to tissue regeneration^7, 8, 67^. It would therefore be interesting to extend our study of the linear rheological properties of hyaluronan hydrogels in the future to the non-linear regime of large deformations and to complex time-dependent effects such as plasticity and self-healing^68^.

## ACKNOWLEDGMENTS

We thank Fred MacKintosh (Rice University, Texas) for useful discussions on polymer rheology and Oleksandr O. Sofronov (AMOLF, The Netherlands) for useful experimental suggestions. This work was funded by the Netherlands Organisation for Scientific Research (NWO, grant number FOM-i41) and was part of the Industrial Partnership Programme Hybrid Soft Materials that is carried out under an agreement between Unilever Research and Development B.V. and NWO.

## ASSOCIATED CONTENT

### Supporting Information

two supplementary methods sections describing the fitting procedure of the transient absorption spectra and the relaxation model; two supplementary figures showing thermoreversibility of the rheology of hyaluronan and the concentration dependence of the flow activation energies.

## For Table of Contents use only

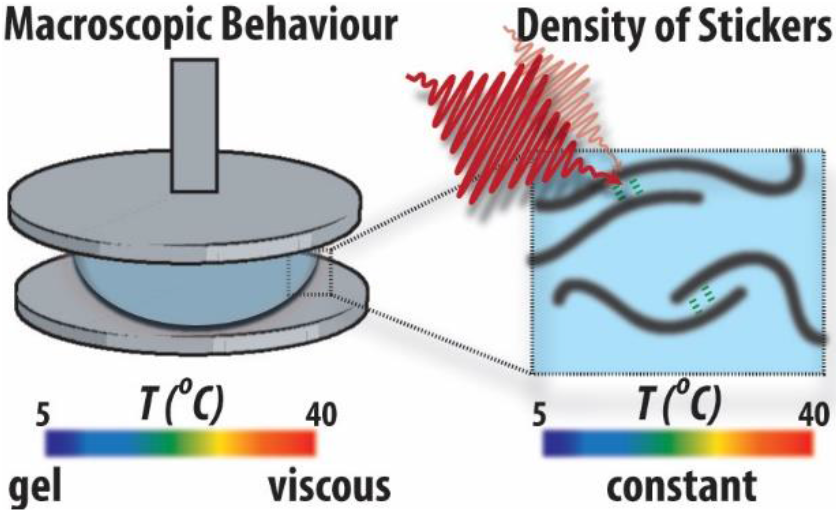

## REFERENCES

(1) Stuart, M. A.; Huck, W. T.; Genzer, J.; Muller, M.; Ober, C.; Stamm, M.; Sukhorukov, G. B.; Szleifer, I.; Tsukruk, V. V.; Urban, M.; Winnik, F.; Zauscher, S.; Luzinov, I.; Minko, S. Emerging applications of stimuli-responsive polymer materials. Nat Mater 2010, 9, 101–13.

(2) Zhang, Z.; Chen, Q.; Colby, R. H. Dynamics of associative polymers. Soft Matter 2018, 14, 2961–2977.

(3) Hofman, A. H.; van Hees, I. A.; Yang, J.; Kamperman, M. Bioinspired Underwater Adhesives by Using the Supramolecular Toolbox. Advanced materials (Deerfield Beach, Fla.) 2018, 30, e1704640.

(4) Li, J.; Mooney, D. J. Designing hydrogels for controlled drug delivery. Nature reviews. Materials 2016, 1.

(5) Rosales, A. M.; Anseth, K. S. The design of reversible hydrogels to capture extracellular matrix dynamics. Nat. Rev. Mater. 2016, 1, 15012.

(6) Stanton, M. M.; Samitier, J.; Sánchez, S. Bioprinting of 3D hydrogels. Lab on a chip 2015, 15, 3111–5.

(7) Highley, C. B.; Prestwich, G. D.; Burdick, J. A. Recent advances in hyaluronic acid hydrogels for biomedical applications. Curr. Opin. Biotechnol. 2016, 40, 35–40.

(8) Wolf, K. J.; Kumar, S. Hyaluronic Acid: Incorporating the Bio into the Material. ACS biomaterials science & engineering 2019, 5, 3753–3765.

(9) Geissler, E.; Hecht, A. M.; Horkay, F. Scaling equations for a biopolymer in salt solution. Phys Rev Lett 2007, 99, 267801.

(10) Berezney, J. P.; Saleh, O. A. Electrostatic effects on the conformation and elasticity of hyaluronic acid, a moderately flexible polyelectrolyte. Macromolecules 2017, 50, 1085–1089.

(11) Lee, J. Y.; Spicer, A. P. Hyaluronan: a multifunctional, megaDalton, stealth molecule. Current opinion in cell biology 2000, 12, 581–6.

(12) Horkay, F.; Basser, P. J.; Londono, D. J.; Hecht, A. M.; Geissler, E. Ions in hyaluronic acid solutions. The Journal of chemical physics 2009, 131, 184902.

(13) Horkay, F.; Magda, J.; Alcoutlabi, M.; Atzet, S.; Zarembinski, T. Structural, mechanical and osmotic properties of injectable hyaluronan-based composite hydrogels. Polymer (Guildf) 2010, 51, 4424–4430.

(14) Cowman, M. K.; Schmidt, T. A.; Raghavan, P.; Stecco, A. Viscoelastic properties of hyaluronan in physiological conditions. F1000Res. 2015, 4, 622.

(15) Zhu, Y.; Granick, S. Biolubrication: hyaluronic acid and the influence on its interfacial viscosity of an antiinflammatory drug. Macromolecules 2003, 36, 973–976.

(16) Shenoy, V.; Rosenblatt, J. Diffusion of macromolecules in collagen and hyaluronic acid, rigid-rod-flexible polymer, composite matrixes. Macromolecules 1995, 28, 8751–8758.

(17) Richter, R. P.; Baranova, N. S.; Day, A. J.; Kwok, J. C. Glycosaminoglycans in extracellular matrix organisation: are concepts from soft matter physics key to understanding the formation of perineuronal nets? Current opinion in structural biology 2018, 50, 65–74.

(18) Kuo, J. C. H.; Gandhi, J. G.; Roseanna, R. N.; Paszek, M. J. Physical biology of the cancer cell glycocalyx. Nature Physics 2018, 14, 658–669.

(19) Neuman, M. G.; Nanau, R. M.; Oruña-Sanchez, L.; Coto, G. Hyaluronic acid and wound healing. Journal of pharmacy & pharmaceutical sciences : a publication of the Canadian Society for Pharmaceutical Sciences, Societe canadienne des sciences pharmaceutiques 2015, 18, 53–60.

(20) Rooney, P.; Kumar, S. Inverse relationship between hyaluronan and collagens in development and angiogenesis. Differentiation; research in biological diversity 1993, 54, 1–9.

(21) Petrey, A. C.; de la Motte, C. A. Hyaluronan, a crucial regulator of inflammation. Frontiers in immunology 2014, 5, 101.

(22) Back, S. A.; Tuohy, T. M.; Chen, H.; Wallingford, N.; Craig, A.; Struve, J.; Luo, N. L.; Banine, F.; Liu, Y.; Chang, A.; Trapp, B. D.; Bebo, B. F., Jr.; Rao, M. S.; Sherman, L. S. Hyaluronan accumulates in demyelinated lesions and inhibits oligodendrocyte progenitor maturation. Nature medicine 2005, 11, 966–72.

(23) Liu, M.; Tolg, C.; Turley, E. Dissecting the Dual Nature of Hyaluronan in the Tumor Microenvironment. Frontiers in immunology 2019, 10, 947.

(24) Rankin, K. S.; Frankel, D. Hyaluronan in cancer - from the naked mole rat to nanoparticle therapy. Soft Matter 2016, 12, 3841–8.

(25) Temple-Wong, M. M.; Ren, S.; Quach, P.; Hansen, B. C.; Chen, A. C.; Hasegawa, A.; D’Lima, D. D.; Koziol, J.; Masuda, K.; Lotz, M. K.; Sah, R. L. Hyaluronan concentration and size distribution in human knee synovial fluid: variations with age and cartilage degeneration. Arthritis research & therapy 2016, 18, 18.

(26) Gribbon, P.; Heng, B. C.; Hardingham, T. E. The molecular basis of the solution properties of hyaluronan investigated by confocal fluorescence recovery after photobleaching. Biophysical journal 1999, 77, 2210–6.

(27) Innes-Gold, S. N.; Pincus, P. A.; Stevens, M. J.; Saleh, O. A. Polyelectrolyte Conformation Controlled by a Trivalent-Rich Ion Jacket. Phys Rev Lett 2019, 123, 187801.

(28) Gibbs, D. A.; Merrill, E. W.; Smith, K. A.; Balazs, E. A. Rheology of hyaluronic acid. Biopolymers 1968, 6, 777–91.

(29) Gatej, I.; Popa, M.; Rinaudo, M. Role of the pH on hyaluronan behavior in aqueous solution. Biomacromolecules 2005, 6, 61–7.

(30) Wu, S.; Ai, L.; Chen, J.; Kang, J.; Cui, S. W. Study of the mechanism of formation of hyaluronan putty at pH 2.5: part I. Experimental measurements. Carbohydrate polymers 2013, 98, 1677–82.

(31) Wu, S.; Ai, L.; Chen, J.; Kang, J.; Cui, S. W. Study of the mechanism of formation of hyaluronan putty at pH 2.5: part II--theoretical analysis. Carbohydrate polymers 2013, 98, 1683–8.

(32) Balazs, E. A.; Cui, J. The story of hyaluronan putty. Bioact. Carbohydr. Diet. Fibre 2013, 2, 143–151.

(33) Giubertoni, G.; Koenderink, G. H.; Bakker, H. J. Direct Observation of Intrachain Hydrogen Bonds in Aqueous Hyaluronan. The journal of physical chemistry. A 2019, 123, 8220–8225.

(34) Leibler, L.; Rubinstein, M.; Colby, R. H. Dynamics of reversible networks. Macromolecules 1991, 24, 4701–4707.

(35) Rubinstein, M.; Semenov, A. N. Dynamics of entangled solutions of associating polymers. Macromolecules 2001, 34, 1058–1068.

(36) Woutersen, S.; Mu, Y.; Stock, G.; Hamm, P. Hydrogen-bond lifetime measured by time-resolved 2D-IR spectroscopy: N-methylacetamide in methanol. Chem. Phys. 2001, 266, 137–147.

(37) Krause, W. E.; Bellomo, E. G.; Colby, R. H. Rheology of sodium hyaluronate under physiological conditions. Biomacromolecules 2001, 2, 65–9.

(38) Charlot, A.; Auzly-Velty, R. Novel hyaluronic acid based supramolecular assemblies stabilized by multivalent specific interactions: rheological behavior in aqueous solution. Macromolecules 2007, 40, 9555–9563.

(39) Giubertoni, G.; Burla, F.; Martinez-Torres, C.; Dutta, B.; Pletikapic, G.; Pelan, E.; Rezus, Y. L. A.; Koenderink, G. H.; Bakker, H. J. Molecular Origin of the Elastic State of Aqueous Hyaluronic Acid. J Phys Chem B 2019, 123, 3043–3049.

(40) Ahmadi, M.; Hawke, L. G. D.; Goldansaz, H.; vanRuymbeke, E. Dynamics of Entangled Linear Supramolecular Chains with Sticky Side Groups: Influence of Hindered Fluctuations. Macromolecules 2015, 48, 7300–7310.

(41) Hackelbusch, S.; Rossow, T.; vanAssenbergh, P.; Seiffert, S. Chain Dynamics in Supramolecular Polymer Networks. Macromolecules 2013, 46, 6273–6286.

(42) Cyphert, J. M.; Trempus, C. S.; Garantziotis, S. Size Matters: Molecular Weight Specificity of Hyaluronan Effects in Cell Biology. International journal of cell biology 2015, 2015, 563818.

(43) Krezel, A.; Bal, W. A formula for correlating pKa values determined in D2O and H2O. Journal of inorganic biochemistry 2004, 98, 161–6.

(44) Lowry, K. M.; Beavers, E. M. Thermal stability of sodium hyaluronate in aqueous solution. Journal of biomedical materials research 1994, 28, 1239–44.

(45) Amunson, K. E.; Kubelka, J. On the temperature dependence of amide I frequencies of peptides in solution. J Phys Chem B 2007, 111, 9993–8.

(46) Selig, O.; Siffels, R.; Rezus, Y. L. Ultrasensitive Ultrafast Vibrational Spectroscopy Employing the Near Field of Gold Nanoantennas. Phys Rev Lett 2015, 114, 233004.

(47) Dea, I. C.; Moorhouse, R.; Rees, D. A.; Arnott, S.; Guss, J. M.; Balazs, E. A. Hyaluronic acid: a novel, double helical molecule. Science 1973, 179, 560–2.

(48) Lodge, T. P.; Rotstein, N. A.; Prager, S. Dynamics of entangled polymer liquids. Do linear chains reptate. Adv. Chem. Phys. 1990, 79, 1–132.

(49) Dodero, A.; Williams, R.; Gagliardi, S.; Vicini, S.; Alloisio, M.; Castellano, M. A micro-rheological and rheological study of biopolymers solutions: Hyaluronic acid. Carbohydrate polymers 2019, 203, 349–355.

(50) Oelschlaeger, C.; Cota Pinto Coelho, M.; Willenbacher, N. Chain flexibility and dynamics of polysaccharide hyaluronan in entangled solutions: a high frequency rheology and diffusing wave spectroscopy study. Biomacromolecules 2013, 14, 3689–96.

(51) Dougherty, R. C. Temperature and pressure dependence of hydrogen bond strength: A perturbation molecular orbital approach. J. Chem. Phys. 1998, 109, 7372.

(52) Tanaka, F.; Edwards, S. F. Viscoelastic properties of physically cross-linked networks. 2. Dynamic mechanical moduli. J. Non-Newtonian Fluid Mech. 1992, 43, 273–288

(53) Ferry, J. D., Viscoelastic properties of polymers. John Wiley & Sons: New York, 1980.

(54) Shamoun, R. F.; Hariri, H. H.; Ghostine, R. A.; Schlenoff, J. B. Thermal transformations in extruded saloplastic polyelectrolyte complexes. Macromolecules 2012, 45, 9759–9767.

(55) Spruijt, E.; Sprakel, J.; Lemmers, M.; Stuart, M. A.; van der Gucht, J. Relaxation dynamics at different time scales in electrostatic complexes: time-salt superposition. Phys Rev Lett 2010, 105, 208301.

(56) Kovács, A.; Nyerges, B.; Izvekov, V. Vibrational analysis of N-acetyl-alpha-D-glucosamine and beta-D-glucuronic acid. J. Phys. Chem. B 2008, 112, 5728–35.

(57) Hamm, P.; Zanni, M., Concepts and methods of 2D infrared spectroscopy. Cambridge University Press: Cambridge, 2011; p 636–636.

(58) Sheu, S.-Y.; Yang, D.-Y.; Selzle, H. L.; Schlag, E. W. Energetics of hydrogen bonds in peptides. Proc. Natl. Acad. Sci. U.S.A. 2003, 100, 12683–12687.

(59) Bolen, D. W.; Rose, G. D. Structure and energetics of the hydrogen-bonded backbone in protein folding. Annu. Rev. Biochem. 2008, 77, 339–362.

(60) Tarek, M.; Tobias, D. J. Role of protein-water hydrogen bond dynamics in the protein dynamical transition. Phys Rev Lett 2002, 88, 138101.

(61) Giannotti, M. I.; Rinaudo, M.; Vancso, G. J. Force spectroscopy of hyaluronan by atomic force microscopy: from hydrogen-bonded networks toward single-chain behavior. Biomacromolecules 2007, 8, 2648–52.

(62) Gerweck, L. E.; Seetharaman, K. Cellular pH gradient in tumor versus normal tissue: potential exploitation for the treatment of cancer. Cancer research 1996, 56, 1194–8.

(63) Corbet, C.; Feron, O. Tumour acidosis: from the passenger to the driver’s seat. Nature reviews. Cancer 2017, 17, 577–593.

(64) Erra Díaz, F.; Ochoa, V.; Merlotti, A.; Dantas, E.; Mazzitelli, I.; Gonzalez Polo, V.; Sabatté, J.; Amigorena, S.; Segura, E.; Geffner, J. Extracellular Acidosis and mTOR Inhibition Drive the Differentiation of Human Monocyte-Derived Dendritic Cells. Cell reports 2020, 31, 107613.

(65) Erra Díaz, F.; Dantas, E.; Geffner, J. Unravelling the Interplay between Extracellular Acidosis and Immune Cells. Mediators of inflammation 2018, 2018, 1218297.

(66) Schmaljohann, D. Thermo- and pH-responsive polymers in drug delivery. Adv Drug Deliv Rev 2006, 58, 1655–70.

(67) Singh, A.; Corvelli, M.; Unterman, S. A.; Wepasnick, K. A.; McDonnell, P.; Elisseeff, J. H. Enhanced lubrication on tissue and biomaterial surfaces through peptide-mediated binding of hyaluronic acid. Nature Materials 2014, 13, 988–995.

(68) Meng, F.; Pritchard, R. H.; Terentjev, E. M. Stress Relaxation, Dynamics, and Plasticity of Transient Polymer Networks. Macromolecules 2016, 49, 2843–2852.

