## Supplementary Information for "Connecting the stimuli-responsive rheology of biopolymer hydrogels to underlying hydrogen-bonding interactions"

#### Supplementary Methods:

**Fit of the transient absorption spectra.** The fitting procedure is described in detail in Ref. [1]. A short summary is provided here. The partial overlap of the –COOH diagonal peak with the cross-peak hinders a straightforward determination of the dynamics of the cross-peak. Specifically, the positive-signed *esa* band of the diagonal peak obscures the negative-signed bleach of the cross-peak. Furthermore, any heating signal originating from the interaction of the ultra-short laser pulses with the sample would either contribute or cancel out bands in 2DIR spectra. Therefore, it is important to deconvolute these contributions in order to extract the dynamics emanating solely due to the cross-peaks.

The transient absorption spectrum following excitation of the C=O vibration is obtained by averaging the signals obtained with pump frequencies between 1700 and 1760 cm$^{-1}$ (C=O pump-region). The transient absorption spectrum following excitation of the C=O mode is expected to be a linear combination of four Lorentzians (two describing the cross-peaks and two describing the diagonal peaks) and a heating spectrum, $h(\omega_j^{probe})$ . As a consequence, our fitting routine is not able to disentangle the contribution of the positive-signed *esa* of the cross-peak. We thus use three Lorentzians, $l_k(\omega_{probe})$, and we describe the transient absorption spectrum:

$$\forall i \text{ for } \omega_{min}^{probe} \leq \omega_j^{probe} \leq \omega_{max}^{probe}$$

$$S\left(\omega_j^{probe}, t_i\right) = \sum_{k=1}^{3} c_k(t_i) l_k\left(\omega_j^{probe}\right) + c_h(t_i) h(\omega_j^{probe}), \quad \text{(S1)}$$

where $c_k$ and $c_h$ represent the amplitudes of the three Lorenzians and of the heating signature, respectively. The heating spectrum is taken directly from the transient absorption spectrum at 10 ps, when the vibrational dynamics is over. The width and center position of the cross-peak and the width of the bleach of the diagonal peak are constrained in a global fit to the transient absorption spectra at all delays following excitation of the C=O mode. For the cross-peak, we set the width equals to 39 $cm^{-1}$ and the center position to 1640 $cm^{-1}$, and for the bleach of the diagonal peak we set the width equals to 38 $cm^{-1}$. Similar values were found in our previous work.[1] The width of the *esa* of the diagonal peak, the anharmonicity (i.e. the shift factor of the *esa* center position with respect to the bleach center position), and the center position of the bleach of the diagonal peak are global parameters in the fit, meaning they are fixed at all delay-times, and only the amplitudes, $c_k$ and $c_h$, are allowed to be different at each delay-time. The global fitting procedure is performed as a function of probe frequency and delay time and it is based on the minimization of the least-square error,

$$\sum_{ij}\left(S\left(\omega_j^{probe}, t_i\right) - S^{exp}\left(\omega_j^{probe}, t_i\right)\right)^2, \quad \text{(S2)}$$

where $S^{exp}$ represents the measured transient absorption spectrum.

**Relaxation model.** The relaxation model was described in detail in Ref.[1]. Briefly, we model the delay-time dependence of the cross-peak of the C=O mode and the AM.I mode (amide I mode) as the result of the relaxation of two carboxylic acid species. One species, with population $N_{th}(t)$, relaxes by energy transfer to lower-frequency modes. These lower-frequency modes are anharmonically coupled to the amide modes. The other carboxylic acid species, with population $N_{ent}(t)$, relaxes via energy transfer to a nearby amide I vibration (*ent*). We thus write the cross-peak transient signal as:

$$\Delta\alpha_{cp}(\omega^{probe}, T_w) = -2\sigma_{AM.I}\ (\omega^{probe})(w(\beta_{AM.I,LFM}) * N_{th}(T_w) + N_{ent}(T_w)) \qquad \text{(S3)}$$

where $\sigma_{AM.I}$ is the amide I cross-section and $w(\beta_{AM.I,LFM})$, a coupling-dependent weighing factor, is introduced to take care of the coupling-dependent energy transfer pathway. Note that $\beta_{AM.I,LFM}$ denotes the vibrational coupling between the amide I and low frequency modes. $N_{th}(T_w)$ and $N_{ent.}(T_w)$ can be described as follows:

$$N_{th}\ (T_w) = N_{th}\ (T_w = O)\left(\frac{T_{1LFM}}{T_{1COOD} - T_{1LFM}}\right)\left(e^{-T_w/T_{1COOD}} - e^{-T_w/T_{1LFM}}\ \right) \qquad \text{(S4)}$$

and

$$N_{ent}(T_w) = N_{ent}\ (T_w = O)\left(\frac{T_{1AM.I}}{T_{ent} - T_{1AM.I}}\right)\left(e^{-T_w/T_{ent}} - e^{-T_w/T_{1AM.I}}\right)$$

(S5)

where $T_{1COOD}$ is the lifetime of the carboxylic acid mode (0.65±0.1 ps), $T_{1AM.I}$ is the lifetime of the amide I mode (0.65±0.1 ps) and $T_{1LFM}$ is the lifetime of the low frequency modes extracted from a fit to the cross-peak signal of the C=O mode and the AM.II mode (amide II mode).[1] By normalizing the $\Delta\alpha_{cp}(\omega^{probe}, T_w)$ to $\Delta\alpha_{COOD}(\omega^{probe}, \overline{T_w} = 200\ fs)$ we obtain from Eq. S3 that

$$\frac{\Delta\alpha_{cp}(\omega^{probe}, T_w)}{\Delta\alpha_{COOD}(\omega^{probe}, \overline{T_w})} = \frac{-2\sigma_{AM.I}\ (\omega^{probe})(w(\beta_{AM.I,LFM}), N_{th}(T_w) + N_{ent}(T_w))}{-2\sigma_{COOD,bl}\ (\omega^{probe})N_{COOD}(\overline{T_w})} =$$

$$= \frac{\sigma_{AM.I}\ (\omega^{probe})}{\sigma_{COOD}\ (\omega^{probe})}\ \frac{(w(\beta_{AM.I,LFM}), N_{th}(T_w) + N_{ent.}(T_w))}{N_{COOD}(\overline{T_w})} = \qquad \text{(S6)}$$

and by using Eq. S4 and Eq. S5 it follows that:

$$= c_{th.} \cdot \left(\frac{T_{1LFM}}{T_{1COOD} - T_{1LFM}}\right)\left(e^{-T_w/T_{1COOD}} - e^{-T_w/T_{1LFM}}\right) + c_{ent.}\left(\frac{T_{1AM.I}}{T_{ent} - T_{1AM.I}}\right)\left(e^{-T_w/T_{ent}} - e^{-T_w/T_{1AM.I}}\right), \tag{S7}$$

Here,

$$c_{th.} \equiv \frac{\sigma_{AM.I}\left(\omega^{probe}\right) w(\beta_{AM.I,LFM}),}{\sigma_{COOD}\left(\omega^{probe}\right)} \frac{N_{th}\left(\overline{T_w}\right)}{N_{COOD}\left(\overline{T_w}\right)} \text{ and } c_{ent.} \equiv \frac{\sigma_{AM.I}\left(\omega^{probe}\right)}{\sigma_{COOD}\left(\omega^{probe}\right)} \frac{N_{ent.}\left(\overline{T_w}\right)}{N_{COOD}\left(\overline{T_w}\right)}$$

are global free parameters of the fit. Because of the limited resolution at long time delay, $c_{th.}$ is constrained to be at least 0.02.[1]

**References**

(1) Giubertoni, G.; Burla, F.; Martinez-Torres, C.; Dutta, B.; Pletikapic, G.; Pelan, E.; Rezus, Y. L. A.; Koenderink, G. H.; Bakker, H. J. Molecular Origin of the Elastic State of Aqueous Hyaluronic Acid. *J Phys Chem B* **2019,** 123, 3043-3049.

### Supplementary figures

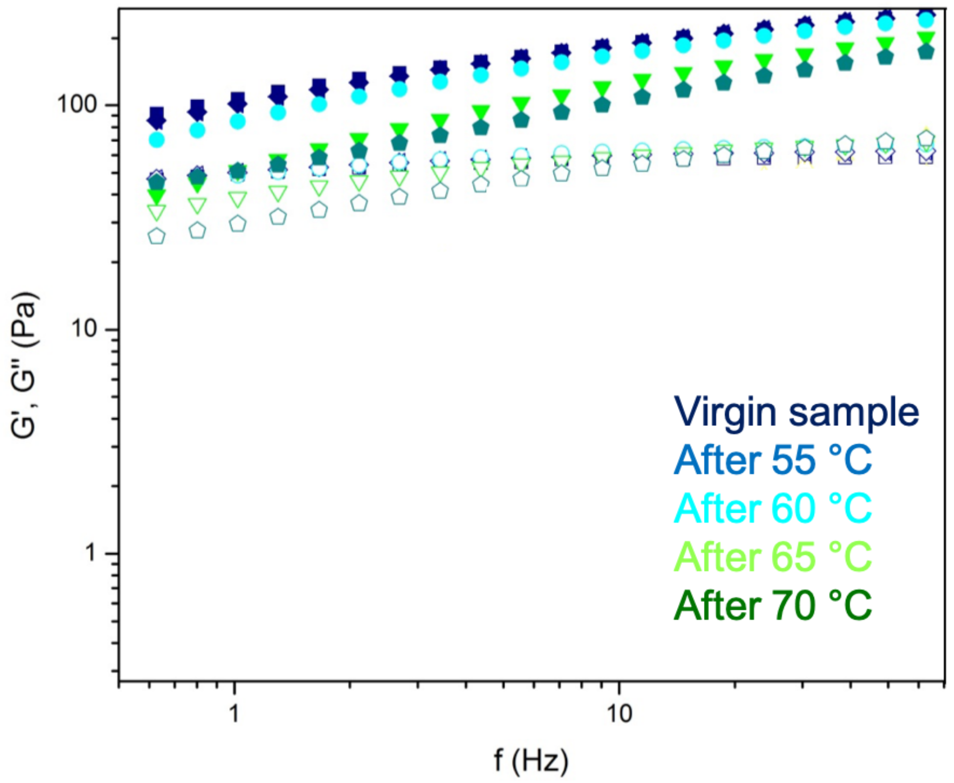


**Supplementary Figure S1: Thermoreversibility of the rheology of hyaluronan solutions at pH 2.5**. Measurements were carried out at a temperature of 20˚C before and after the sample was heated to 55˚C, 60˚C, 65˚C or 70 ˚C. Both the elastic modulus *G*' (solid circles) and the viscous modulus *G*'' (empty circles) return back to the original moduli at 20˚C as long as the temperature does not exceed 60˚C.

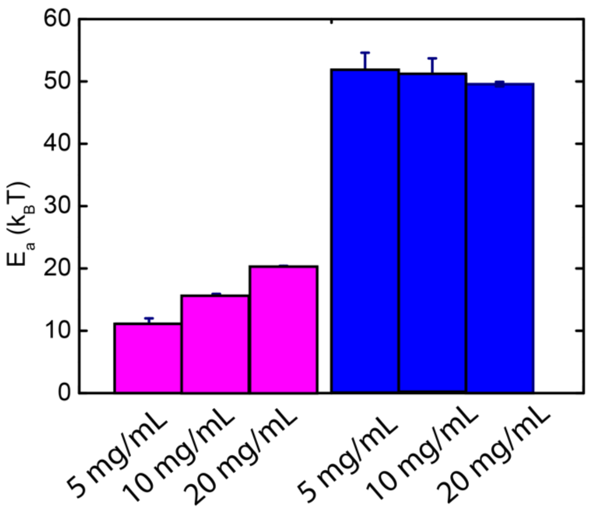


**Supplementary Figure S2:** Concentration dependence of the flow activation energies for hyaluronan solutions at pH 7.0 (pink) and at pH 2.5 (blue). The concentration was varied from 5 to 20 mg/mL, as labelled.
